# Decoupling Topology from Geometry: Detecting Large-Scale Conformational Changes via Conformational Scanning

**DOI:** 10.64898/2026.03.28.714756

**Authors:** Runfeng Lin, Sebastian E. Ahnert

## Abstract

Protein function is fundamentally driven by structural dynamics, yet the majority of structural bioinformatics treats proteins as static rigid bodies. While Molecular Dynamics (MD) simulations attempt to capture these motions, they are computationally prohibitive for exploring large-scale conformational changes, such as domain movements or allostery, which occur on timescales often inaccessible to standard simulation. However, the Protein Data Bank (PDB) contains a latent wealth of dynamic information in the form of redundant entries proteins solved in multiple distinct conformational states. Detecting these “shape-shifting” pairs remains challenging because standard structural alignment algorithms (e.g., TM-align) rely on rigid-body superposition, which fails when substantial geometric rearrangement occurs. In this study, we introduce a high-throughput method to systematically mine the PDB for proteins that share identical topology but exhibit divergent tertiary conformations. By utilizing a coarse-grained Secondary Structure Element (SSE) representation, we decouple topological connectivity from geometric rigidity, allowing for the detection of conformational homologues that share low global structural similarity despite high predicted structural similarity. We applied this “conformational scanning” across the entire RCSB database, identifying a curated dataset of proteins undergoing significant structural rearrangements. This work bridges the gap between static structural data and dynamic function, providing a critical “ground truth” dataset for benchmarking data-driven protein design and checking the plausibility of generative structure models.

## 1 Introduction

Protein structures play a crucial role in determining biological function, yet they are not static entities; rather, they can adapt into multiple distinct conformations in response to interacting ligands or environmental changes. Consequently, proteins should be viewed as dynamic ensembles governed by complex energy landscapes rather than as single static states [1, 2]. For instance, essential processes such as enzymatic catalysis and signal transduction rely on the ability to access specific “excited” states or cryptic conformations [3]. Therefore, characterizing the dynamics of these structures is essential for understanding biological mechanisms like induced-fit binding and allostery. Remarkably, the trigger for these conformational changes is often simple: the binding of a single small molecule can drive the large-scale rearrangement of a macromolecule composed of thousands of atoms. Beyond fundamental biology, the application of these mechanics is broad; the ability to engineer conformational switches is currently driving the development of next-generation biosensors and smart biomaterials that react to their environment [4].

Early studies of protein conformations heavily utilized Molecular Dynamics (MD) simulations. This well-established paradigm provides atomic-level details of structural properties within a given timescale, thereby enabling the study of protein dynamics and conformational shifts [5, 6]. However, as proteins are inherently allosteric and dynamic, using direct MD simulations to sample transitions between distinct conformations requires a significant amount of computing resources [3]. To overcome this bottleneck, various methods have been developed to accelerate the sampling process. An early approach is Locally Enhanced Sampling (LES), which facilitates the crossing of energy barriers by modelling specific flexible regions with multiple non-interacting copies [7, 8]. Other notable techniques include Grow-to-Fit Molecular Dynamics (G2FMD), designed for *ab initio* side-chain refinement to prevent atoms from becoming trapped in high-energy steric clashes [9], and Coarse-Grained or Elastic Network Models (ENM), which simplify the protein representation to capture essential dynamics at a fraction of the computational cost [10, 11].

The recent advent of Machine Learning (ML) models has enabled the accurate prediction of atomic details in protein structures. Building on this progress, significant efforts have been made to predict conformational changes using ML approaches. To address the scarcity of training data, Hu et al. simulated the transitions of proteins known to exhibit multiple states. These data were then utilized to train an ML model capable of predicting specific high-energy transition states and the non-linear pathways connecting two stable conformers [12]. Complementarily, Liu et al. introduced a framework leveraging out-of-distribution detection in hyperspherical latent spaces to automatically pinpoint high-energy transition states from simulation data, which often remain elusive to traditional clustering methods [13].

While these computational approaches provide valuable insights, they often rely on synthetic data or extensive sampling. In this work, we propose an alternative data-driven strategy: analysing potential conformational transitions by decoupling topological similarity from geometric alignment. Proteins can present high topological similarity despite low sequence identity [14], making it difficult to detect conformational relationships using standard sequence or rigid-structure alignment, especially when domain movements result in a low TM-score [15, 16]. To systematically mine the database for these “cryptic” conformational changes, we adapted the Secondary Structure Elements (SSEs) framework [17]. This framework has previously been used to show that the connectivity of SSEs robustly reflects protein structural properties and folding [18, 19]. Here, protein pairs that exhibit a high *predicted* TM-score (based on SSE topology) but a low *actual* TM-score (based on geometry) are identified as candidates for conformational change. By utilizing a dual-conformational scanner (illustrated in Figure 1), we demonstrate that the TM-scores of these pairs can be significantly improved by identifying maximally similar subregions of the two proteins in a given pair, and thereby confirm the likely presence of conformational change.

**Figure 1:**
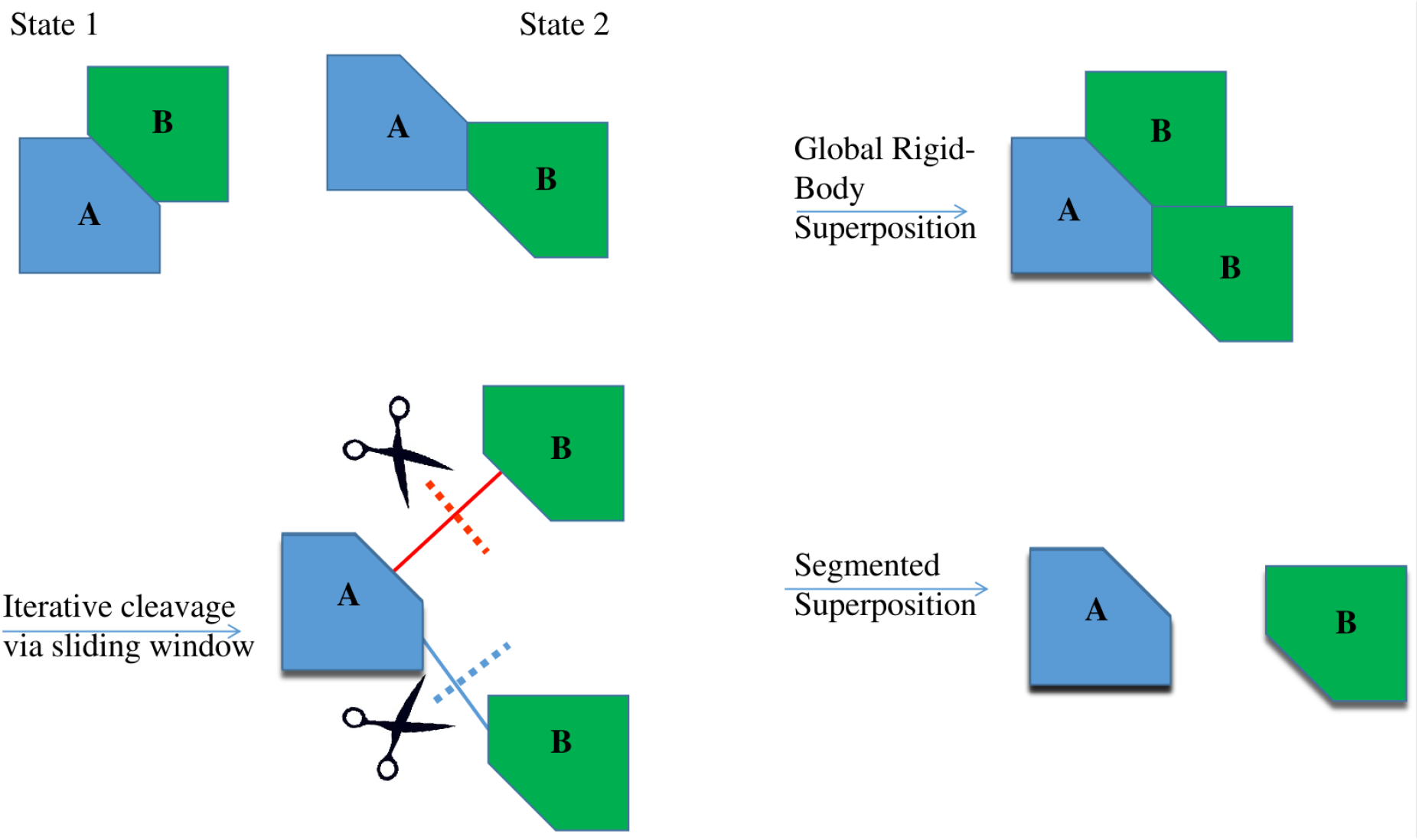
Schematic of the dual-conformational scanning pipeline. Traditional global superposition (top path) fails to capture true structural similarity when large-scale domain movements occur, resulting in a poor overall alignment. To address this, our method iterates a cleavage point across the alignment using a sliding window to split the structure into independent rigid bodies (bottom path). These sub-structures are then superimposed independently, accurately reflecting the conserved topological elements despite spatial divergence.

## 2 Results

### 2.1 Global Detection of Conformational Changes in the RCSB Database

To systematically survey the landscape of protein conformational plasticity, we extended our analysis to the entire RCSB database [20]. After excluding sequences with high content of unknown residues and chains containing fewer than five Secondary Structure Elements (SSEs), the curated dataset comprised 548,782 sequences.

While the SSE-based alignment is computationally efficient, a comprehensive all-to-all comparison of this dataset (*N ≈* 5.5 × 10^5^) would generate an unmanageable volume of alignment data. To address this, we employed the heuristic filter described by Lin et al. [17], which significantly accelerates the search process by reducing the requisite number of alignments without compromising sensitivity.

We established a rigorous criterion to identify “cryptic” conformational changes: pairs must exhibit high topological similarity (Predicted TM-score > 0.5) but significant geometric divergence (Actual TM-score < Predicted TM-score ™0.1). This differential threshold isolates pairs where the connectivity of secondary structures remains conserved despite large-scale spatial rearrangements. This screening protocol identified approximately 146 million pairs exhibiting a discrepancy of > 0.1 between predicted and actual TM-scores.

These candidate pairs were subsequently analysed using the conformational scanner to determine whether their geometric divergence could be reconciled through rigid-body domain adjustments. The results reveal a widespread prevalence of topological robustness: 81% (115 million) of the identified candidates exhibited a TM-score improvement of > 0.01, and 74 million achieved an improvement of at least 0.1 following the 2-fold conformational adjustment (Figure 3). As illustrated in Figure 2, the distribution of TM-scores shifts significantly post-adjustment. The Kolmogorov-Smirnov (KS) test confirms this shift (Figure 2c), yielding a maximum distance (*D*) of 0.529 between the cumulative distribution functions of the original and adjusted states. These findings demonstrate that our topological mining approach effectively uncovers significant conformational metamorphosis—geometric transitions that are substantial enough to elude standard rigid-structure alignment yet are ubiquitous in the structural archive.

**Figure 2:**
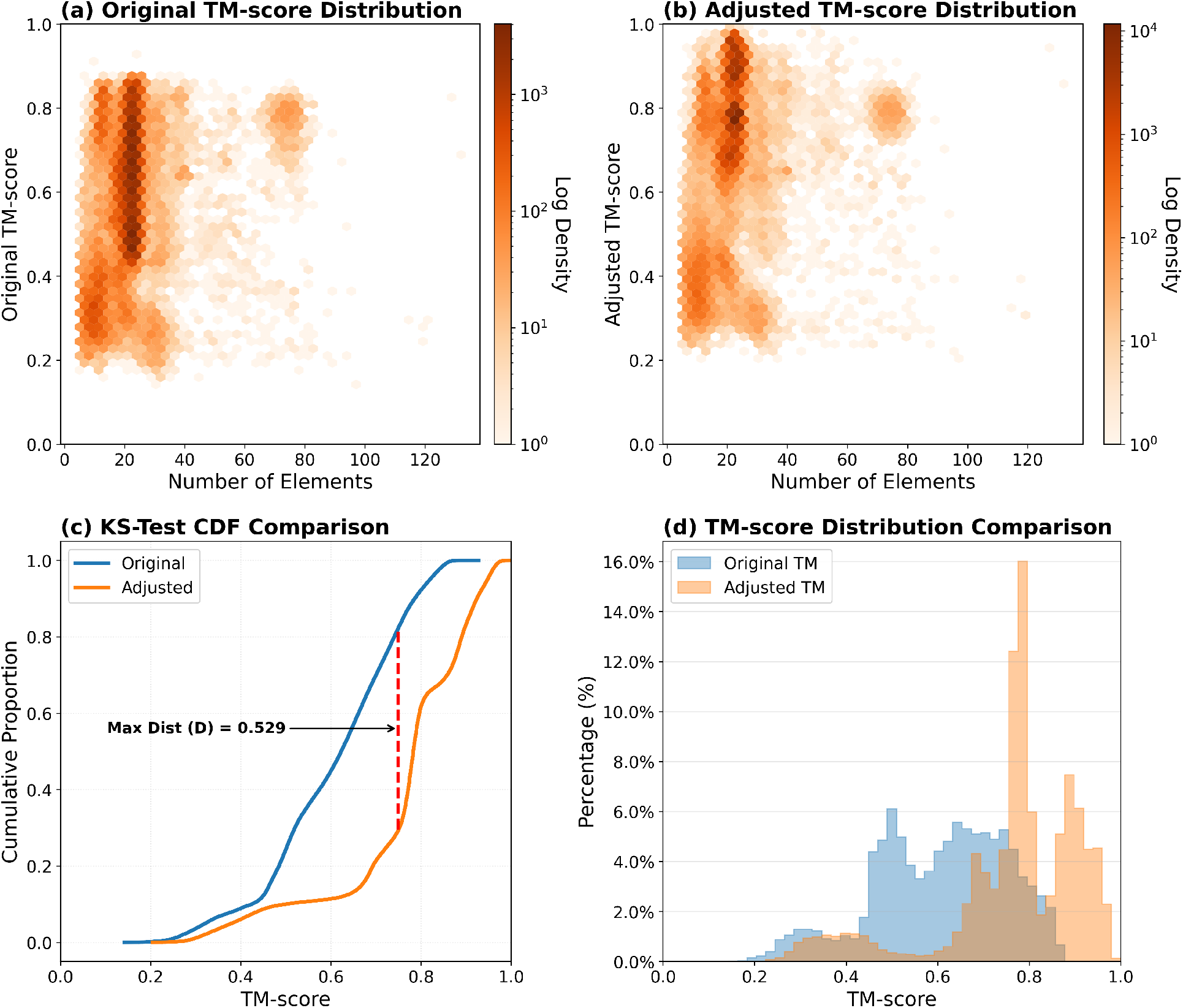
Global Analysis of Conformational Plasticity Across the RCSB Database. (a) Density distribution of original TM-scores versus the number of secondary structure elements (SSEs) for candidate pairs, showing a broad spread of structural similarity. (b) Density distribution of adjusted TM-scores, revealing a distinct shift toward higher structural similarity (darker density at high TM-scores) after accounting for domain movements. (c) Cumulative Distribution Function (CDF) comparison between Original (blue) and Adjusted (orange) TM-scores. The large maximum distance (*D* = 0.529) quantifies the significant recovery of structural similarity. (d) Histogram comparison illustrating the population shift; the adjusted distribution (orange) is markedly skewed toward the high-similarity regime compared to the original geometric alignment (blue).

**Figure 3:**
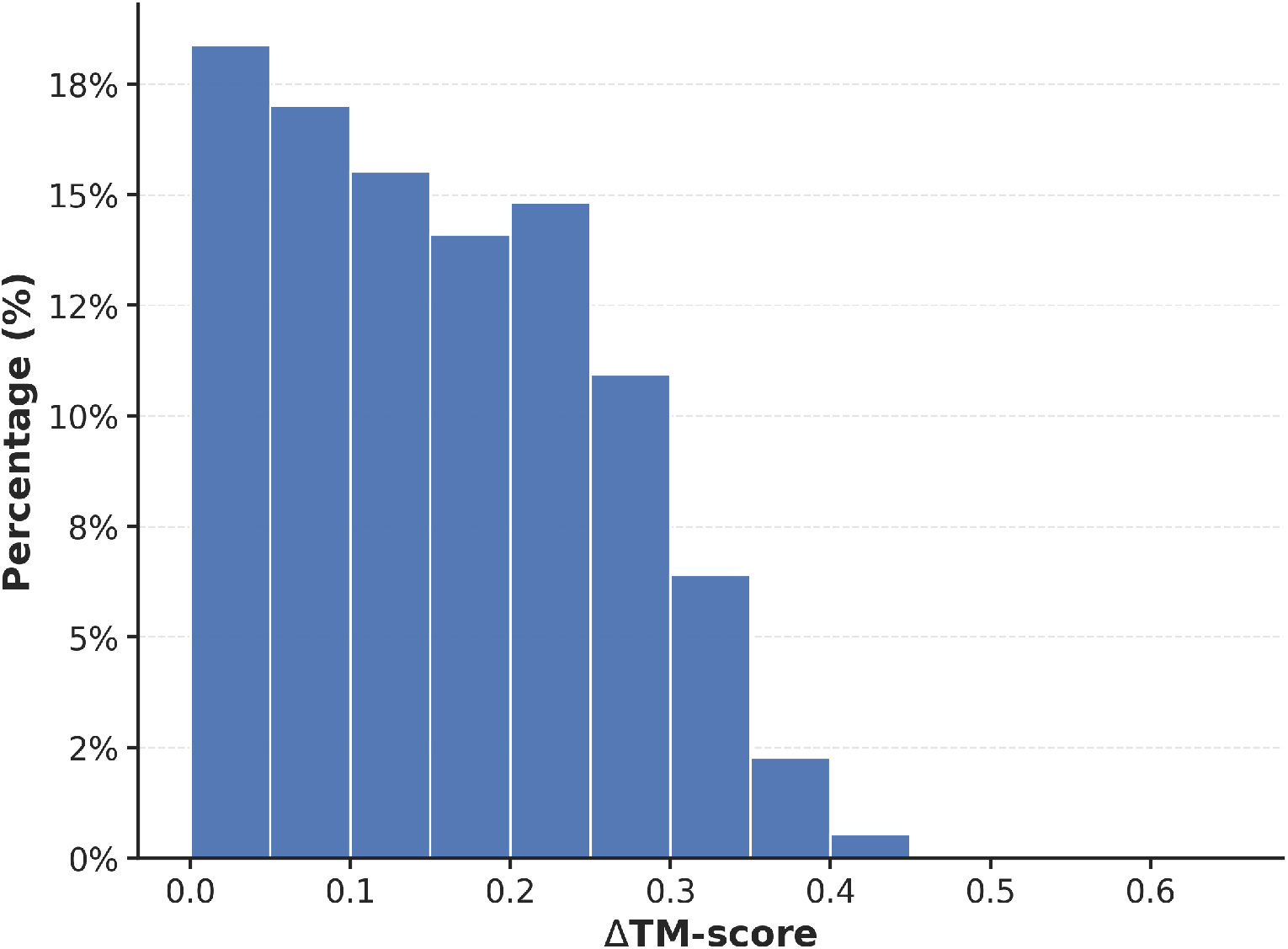
Distribution of TM-score improvement (ΔTM-score) following conformational scanning. The histogram details the percentage of candidate pairs exhibiting various degrees of structural alignment improvement after a 2-fold rigid-body domain adjustment. The right side of the histogram shows that millions of protein pairs get a major boost in their TM-score (ΔTM-score ≥ 0.1) once we account for domain movements.

### 2.2 Decoupling Sequence Identity from Structural Similarity

To assess whether our conformational scanner captures structural relationships that are invisible to sequence-based methods, we analysed the correlation between sequence identity and TM-score before and after adjustment. A major challenge in structural biology is the “Twilight Zone” of homology, where sequence identity falls below 0.3, making evolutionary relationships difficult to detect [14]. Our dataset contains over 88.3 million such pairs (comprising 469,845 unique proteins). For this challenging subset, the mean structural improvement (ΔTM) following adjustment is 0.134, indicating a significant recovery of geometric matching.

Crucially, our method “rescues” a vast number of potential conformational isomers that would be discarded by standard alignment. Among the pairs in the Twilight Zone, over 73 million achieve an adjusted TM-score > 0.5, a threshold widely accepted as the indicator of a shared fold [21]. The mean improvement for this rescued subset is even higher at 0.151. As shown in Figure 4(d), the population distribution shifts dramatically: while the original geometric alignment (blue) suggests these pairs are unrelated, the adjusted distribution (orange) reveals a hidden layer of structural conservation. Notably, approximately 81% of the total ‘Twilight Zone’ pairs exceeds a TM-score of 0.6 after adjustment—a threshold where the probability of sharing the same topology rises to over 90% [21].

**Figure 4:**
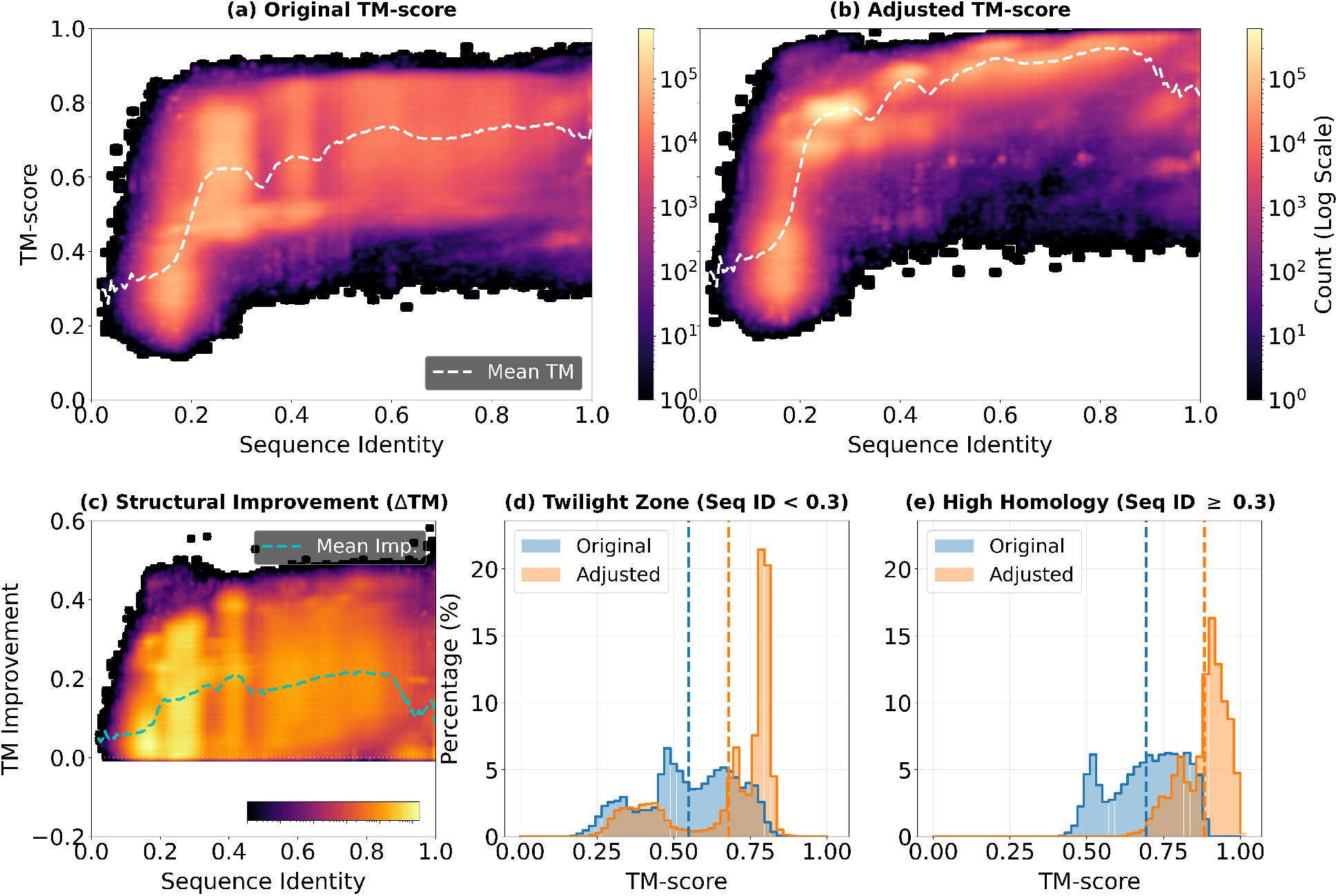
Decoupling Topological Similarity from Sequence Identity. (a) Density plot of Original TM-score vs. Sequence Identity, showing weak structural correlation in the low-identity regime. (b) Density plot of Adjusted TM-score vs. Sequence Identity, revealing a recovered signal of structural similarity (high TM-score) even for sequences in the “Twilight Zone” (Identity < 0.3). Dashed lines in (a) and (b) represent the conditional mean TM-score. (c) Structural Improvement (ΔTM) as a function of sequence identity; the method provides the largest geometric corrections for remotely related sequences. (d) Comparative histograms for the “Twilight Zone” subset (Identity < 0.3). The shift from the original (blue) to adjusted (orange) distribution illustrates the “rescue” of millions of evolutionary pairs into the structurally significant regime. (e) Comparative histograms for the High Homology subset (Identity ≥ 0.3). The adjusted distribution (orange) shows near-universal structural similarity, effectively eliminating pairs that artificially fell below TM-score < 0.5 during rigid-body alignment. The contrast between (d) and (e) demonstrates the scanner’s ability to detect both deep evolutionary relationships and highly dynamic conformational changes.

Furthermore, this scanner is highly effective at identifying conformational changes within the High Homology regime, where structural differences are primarily driven by dynamics. As shown in Figure 4(e), a significant number of homologous pairs failed to be detected by standard rigid-body alignment, yielding TM-scores < 0.5. After adjustment, these scores recover dramatically, suggesting that this approach accurately resolves both deep evolutionary homology and large-scale dynamic flexibility.

### 2.3 Validation against CATH structural classification

To further validate whether these conformationally distinct and sequentially diverse proteins hold biological significance, we mapped the identified pairs to the CATH structural classification database [22]. Focusing on the “Twilight Zone” subset (sequence identity < 0.3), we identified approximately 14.5 million pairs where both proteins possess CATH domain annotations. Strikingly, 14.2 million of these pairs (98.0%) were found to share the same CATH superfamily. This high recovery rate suggests that our method is not only robust to large-scale conformational changes but also biologically accurate in detecting distant evolutionary relationships that sequence alignment misses.

However, as illustrated in Figure 5, the distribution of these families is highly skewed—a known characteristic of the PDB. Approximately 13.1 million pairs belong to a single hyper-abundant superfamily (CATH 2.60.40.10). Even after excluding this dominant family to control for data bias, we found that 82% of the remaining pairs still share the same CATH topology. This confirms that our approach effectively captures structural conservation across a diverse range of protein folds, rather than being an artifact of a single abundant topology.

**Figure 5:**
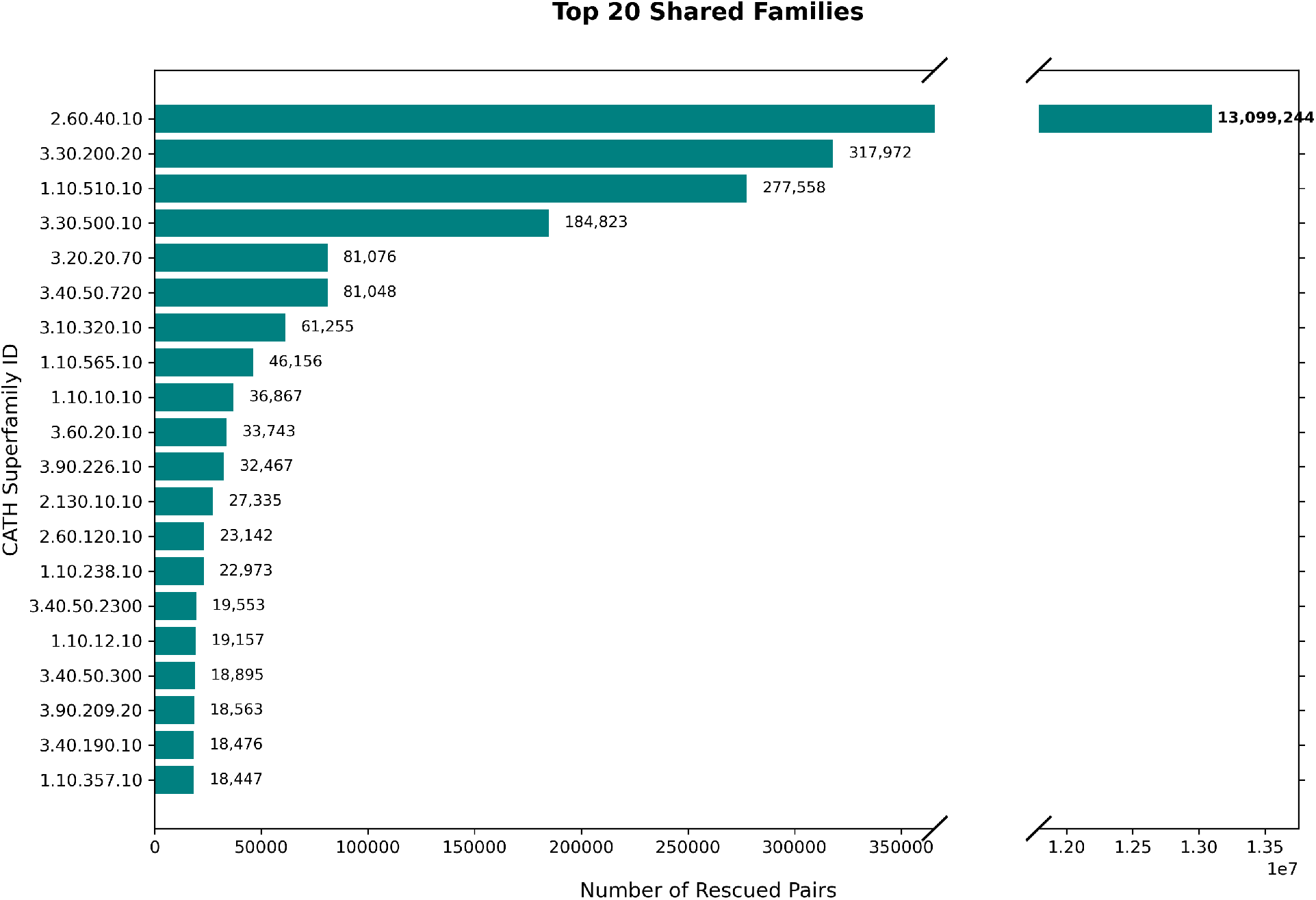
Distribution of CATH superfamilies among Twilight Zone protein pairs. The dataset is heavily dominated by the Immunoglobulin-like fold (CATH 2.60.40.10), which accounts for approximately 90% of the structurally similar pairs. The plot illustrates the top 20 most abundant superfamilies; despite the skew, the method recovers shared topologies across diverse structural classes.

To investigate the extent of structural divergence among the identified pairs, we analysed the subset of “cross-family” matches—pairs that share high structural similarity (Adjusted TM-score > 0.5) but are classified into different CATH superfamilies. While this distribution remains skewed, it is notably less dominated by a single topology compared to the shared-family dataset; the most abundant cross-family connection accounts for only 30% of the total, suggesting a broader diversity of structural bridges.

Figure 6(a) dissects these mismatches by their depth within the CATH hierarchy. Contrary to the expectation that most mismatches would occur at the subtle Topology level (Level 3), we found this category to be the least common (8.7%). Instead, the data is dominated by mismatches at the **Architecture level (Level 4)**, which account for 36.4% of all cross-family pairs. This surprising prevalence indicates that our SSE-based alignment captures a vast network of proteins that share similar topologies but regarded as different homologies which might be slightly functional similar. Although the percentage of mismatch in different class and different architectures are close to different

**Figure 6:**
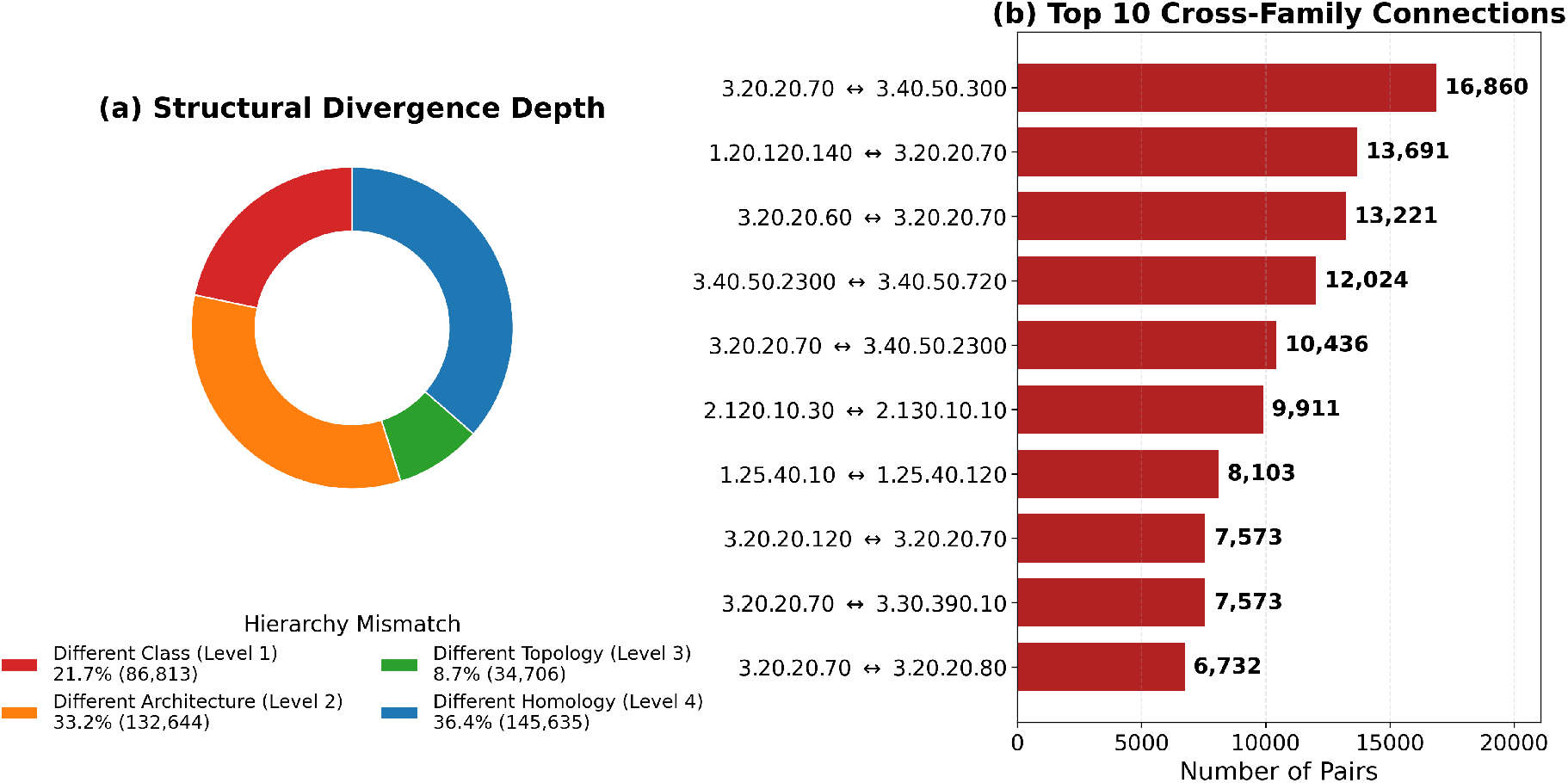
Structural divergence of cross-family Twilight Zone pairs. (a) Hierarchy depth of structural mismatches. The distribution is dominated by Level 4 (Homology) mismatches, indicating that many identified pairs bridge similar topologies that are functionally various. (b) Top 20 specific cross-family connections. The labels denote the CATH superfamilies being bridged (e.g., Family A ↔ Family B).

To determine whether the magnitude of structural correction depends on evolutionary closeness, we compared the TM-score improvement (ΔTM) for pairs within the same CATH superfamily (“same-family”) versus those bridging different families (“cross-family”). As shown in Figure 7a, while same-family pairs exhibit a higher mean improvement (0.153) and a longer tail of extreme improvements (with some ΔTM > 0.5), cross-family pairs still demonstrate a notable mean improvement (0.053). The fact that cross-family pairs—which likely share similar overall topologies despite distant homology—still yield meaningful improvements demonstrates that our SSE-based alignment is a global and generalizable method. It is not biased exclusively toward easily alignable homologues; rather, it is highly effective at resolving large-scale geometric discrepancies in remotely related or structurally divergent proteins that defy standard sequence-based classification.

**Figure 7:**
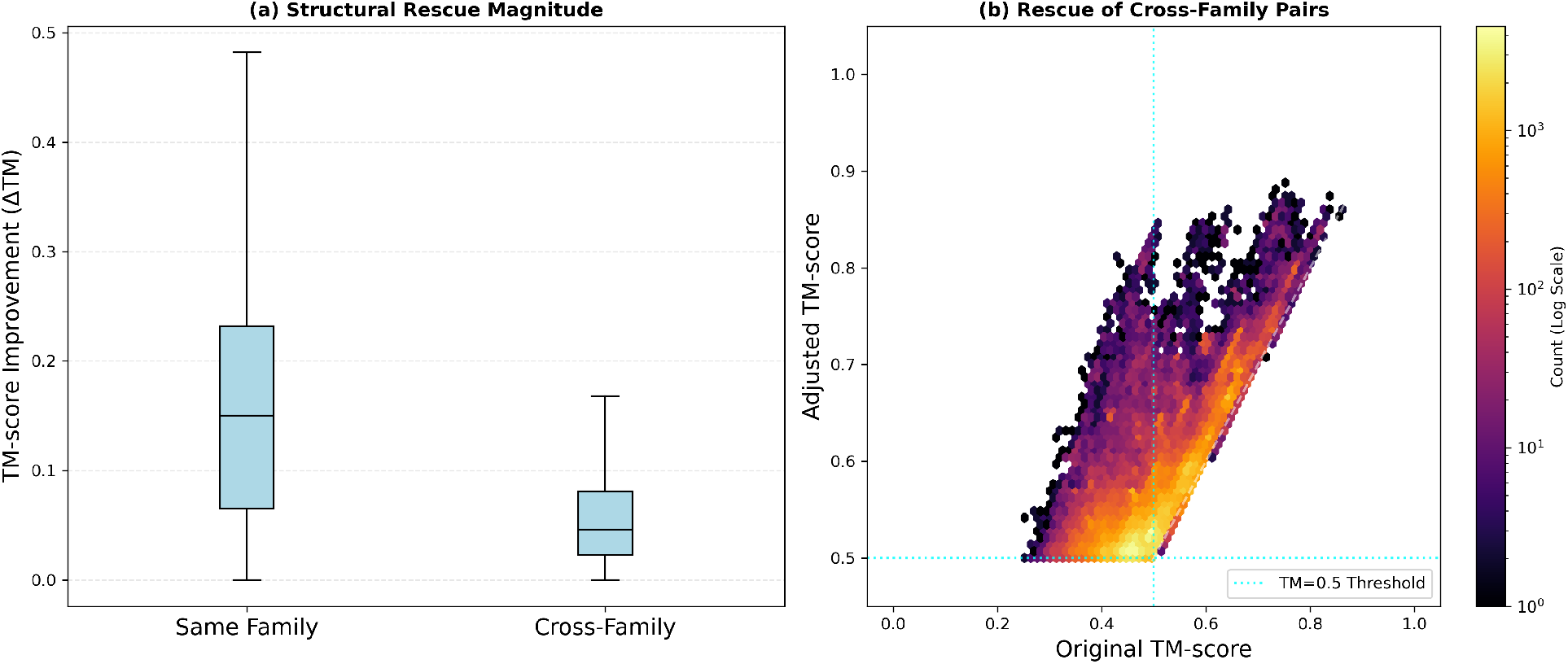
Comparative analysis of structural improvement (ΔTM). (a) Distribution of TM-score improvement for same-family versus cross-family pairs. Although same-family pairs exhibit a higher mean improvement (0.153) and a wider range of extreme improvements (ΔTM > 0.5), the improvement in cross-family groups remains substantial (mean = 0.053), indicating robust performance across different evolutionary distances. (b) Density scatter plot correlating the original and adjusted TM-scores for cross-family pairs, highlighting the population shifted from low similarity (Original TM-score < 0.5) to structural significance (Adjusted TM-score > 0.5).

Figure 8 presents a representative case study of a cross-family protein pair (PDB: 4WV3 B and 5U95 C) that exemplifies the challenge of flexible alignment. Despite sharing a low sequence identity of 25.1%, these proteins exhibit remarkably high structural similarity when adjusted for flexibility (Adjusted TM-score = 0.888), a significant improvement over the standard rigid-body alignment (TM-score = 0.748). Structurally, these proteins consist of two distinct compact domains connected by flexible linker regions. By applying our dual-conformational scanner, the structures were automatically segmented into two major rigid bodies, enabling the successful and independent alignment of both domains (Figure 8b, c).

**Figure 8:**
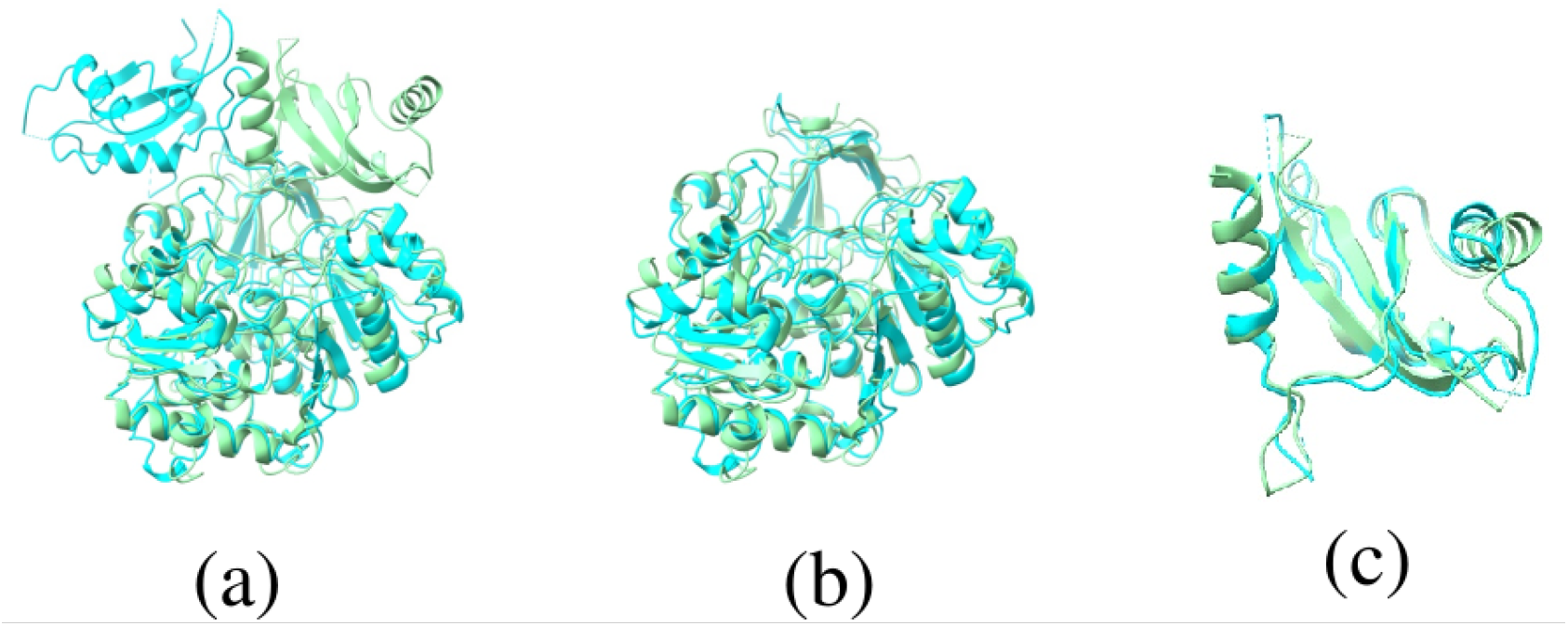
Structural plasticity in the cross-family pair 4WV3 B and 5U95 C. (a) Standard rigid-body alignment (TM-score = 0.748), highlighting the significant spatial displacement between the light green (4WV3 B) and cyan (5U95 C) structures due to inter-domain flexibility. (b, c) Automated dual-conformational alignment (Adjusted TM-score = 0.888, closely matching the predicted TM-score of 0.891). Our scanner successfully identifies and independently aligns the two distinct rigid bodies, shown here as the primary larger domain (b) and the secondary smaller domain (c).

### 2.4 Conformational Clusterings

Although pairwise conformational scanning unravels many pairs of proteins that are in different conformational states, it might introduce redundant results, as one protein might be a conformational homologue to many others.

To reduce this redundancy, the identified pairs were clustered into non-redundant groups: evolutionary clusters, where proteins are linked by sequence identities < 0.3, and dynamic clusters, where proteins are connected by sequence identities ≥ 0.3. According to Figure 9(a), the distribution is highly skewed—a characteristic frequently found in Genotype-Phenotype distributions [23]—and follows a power-law distribution. Specifically, most protein folds are represented by only a few conformational states, while a small subset contains hundreds of distinct entries. In addition, these distributions are almost identical for both evolutionary and dynamic pairs, which suggests that conformational flexibility is a fundamental property of protein folding that is independent of sequence conservation.

**Figure 9:**
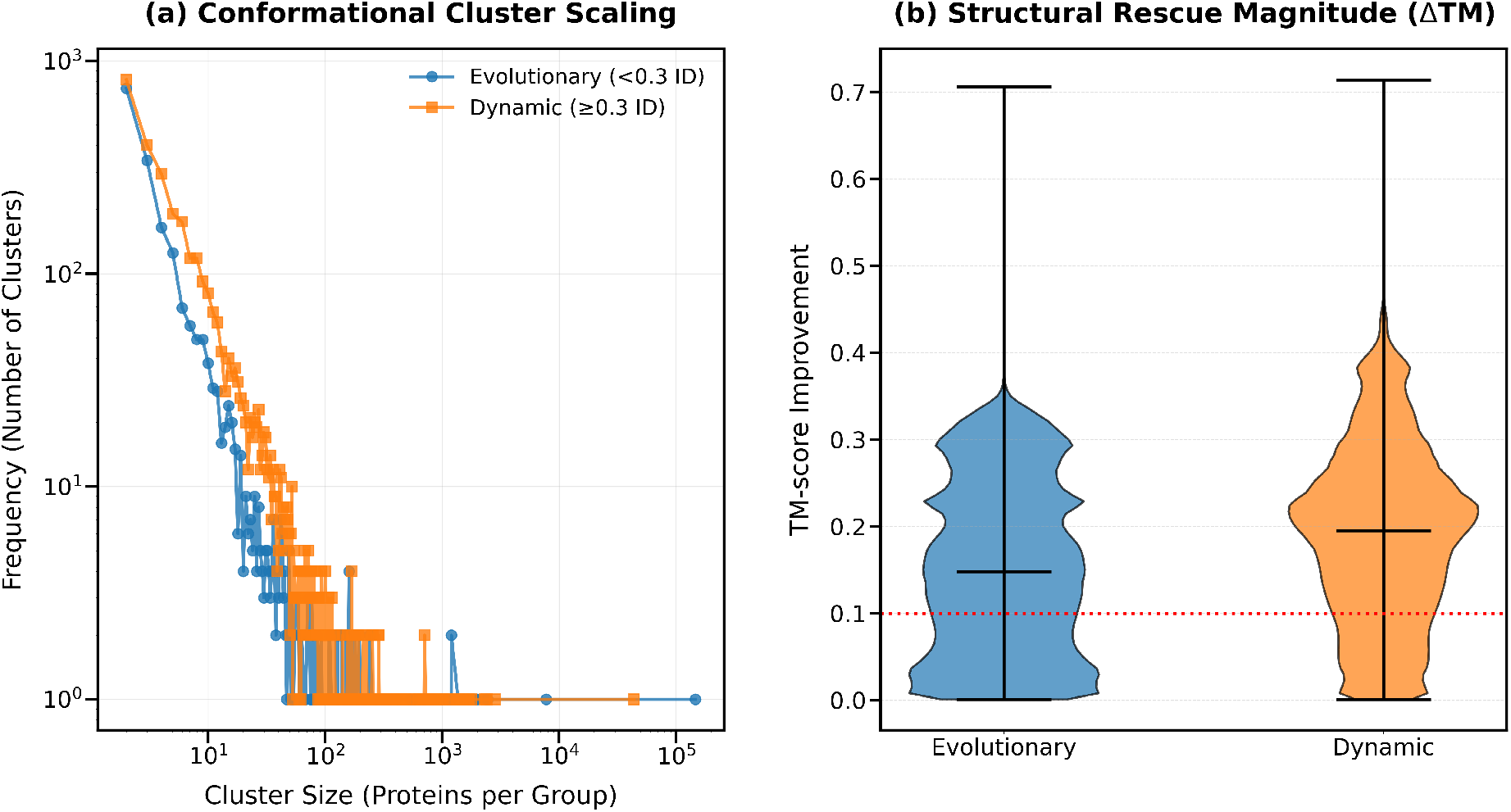
Global scaling and structural rescue magnitude of conformational clusters. (a) Scaling behaviour of conformational cluster sizes for evolutionary (Sequence ID < 0.3) and dynamic (Sequence ID ≥ 0.3) datasets. The log-log plot reveals a power-law distribution where a few highly abundant structural “hubs” contain the majority of conformational entries. (b) Violin plots illustrating the distribution of TM-score improvements (ΔTM) following conformational adjustment. The horizontal black bars denote the mean improvement for each group, both of which comfortably exceed the 0.1 threshold (red dotted line) which we used as a criterion to define significant structural rescue.

Figure 9(b) displays the TM-score improvements (ΔTM) following adjustment. Both cluster types exhibit mean improvements exceeding the 0.1 threshold for significant conformational change. Furthermore, the cluster-based mean improvements (0.154 for evolutionary clusters and 0.191 for dynamic clusters) reveal an interesting contrast when compared to the pairwise-based averages (0.134 for the twilight zone and 0.190 for high homology). This disparity suggests that for evolutionary flexibility, smaller clusters tend to exhibit higher structural flexibility than more abundant ones, whereas no significant size-dependent difference is observed for dynamic flexibility.

## 3 Discussion and Conclusion

In this work, a conformational scanner was implemented across the RCSB Protein Data Bank [20] to generate a comprehensive database of conformationally flexible proteins. This scanner, adapted from our previous SSE alignment framework [17], focuses specifically on the discrepancy between topological connectivity (predicted structural similarity) and rigid-body geometry (ground truth alignment) [15]. Upon systematic identification, we observed that many protein pairs share similar SSE topologies but exhibit drastically divergent geometric conformations. Our dual-conformational scanner effectively resolves these differences, successfully capturing large-scale domain movements. Furthermore, we demonstrated that this pipeline not only recovers conformational changes within highly conserved sequences, but also identifies flexible proteins in the “Twilight Zone” of sequence identity, thereby serving as a powerful complement to traditional sequence-based methods.

Beyond the simple validation of improved structural similarity, we cross-referenced our results with the CATH database [22] for protein pairs located within the “Twilight Zone”. The remarkably high agreement between the established CATH classification and our structural recoveries suggests that our method is not merely identifying structural artifacts, but rather capturing evolutionary relationships that hold true biological meaning.

In addition, current protein structure prediction models primarily output static conformations and lack a deep understanding of protein conformational flexibility [24]. To address this gap, we provide a comprehensive database of structures undergoing significant conformational changes. This curated dataset can serve as a critical benchmark for the future development of machine learning models, ultimately facilitating protein structure prediction as dynamic conformational ensembles.

Although our method has successfully identified a large number of conformationally flexible proteins, it currently operates as a dual-conformational scanner that splits proteins into only two rigid bodies. Consequently, it lacks the ability to accurately capture the conformational flexibility of proteins undergoing more complex shifts involving three or more independent domains. A generalisation of this approach to more than two domains is likely to yield further examples of conformational changes, including more complex rearrangements of larger, multi-domain proteins.

## 4 Method

### 4.1 Definition and Extraction of Secondary Structure Elements (SSEs)

In this study, Secondary Structure Elements (SSEs) are defined based on the continuous sequence length of assigned structural motifs. Initial secondary structure annotations are derived directly from the atomic coordinates of the proteins using the DSSP algorithm under its scheme-2 criteria [25]. To translate these atomic-level annotations into a discrete structural sequence, we extract the lengths of uninterrupted periodic elements (*α*-helices and *β*-strands), while explicitly disregarding unstructured random coils.

To effectively manage the high variance in segment lengths and avoid generating an overly sparse vocabulary, we implemented a variable-resolution tokenization scheme:

- **High-Resolution Encoding:** For shorter structural segments ranging from 2 to 10 residues, each specific length is assigned a distinct alphabetical character (for instance, a length of 2 is denoted as ‘A’, and a length of 3 as ‘B’).
- **Binned Encoding:** To mitigate vocabulary sparsity caused by unusually long elements, extended segments are grouped into discrete bins. Lengths between 12 and 30 residues are grouped with a step size of 2, and any segment exceeding 30 residues is categorized using a step size of 3.

To preserve the topological identity of the protein fold, the structural type of each SSE is differentiated solely through alphabetical case. Specifically, *β*-strands are represented by uppercase letters (e.g., three consecutive strand residues `EEE` map to `‘B’`), whereas *α*-helices are denoted by their lowercase counterparts (e.g., `HHH` maps to `‘b’`). The full source code for this structural extraction and translation pipeline can be accessed via our associated GitHub repository [17].

### 4.2 Heuristic K-mer Filtration

To circumvent the computational bottleneck of an exhaustive all-to-all alignment, the heuristic filtration strategy introduced in [17] was employed to drastically reduce the search space. Specifically, this algorithm identifies highly probable conformational candidates through a highly parallelized, two-stage sequence matching process.

The filtration pipeline leverages a dual-indexing system stored in a high-performance key-value database (RocksDB) [26]. It queries both exact *k*-mers (default *k* = 3) via a sliding window and spaced seeds (matching alternating positions, e.g., indices *i, i* + 2, *i* + 4) to capture both continuous and gapped structural similarities. To distinguish true alignments from random noise, the algorithm tracks the relative positional offsets (diagonal hits) between the query and target sequences.

A target sequence is only retained as a valid candidate if it passes a rigorous two-stage heuristic filter:

1. **Diagonal Consistency:** The sequence must accumulate a minimum number of hits along a single geometric diagonal (scaling with the query length), ensuring contiguous structural conservation.
2. **Aggregate Scoring:** The total number of valid hits across the sequence must exceed a secondary, length-dependent global threshold.

By utilizing persistent database connections and a multi-worker parallel architecture, this heuristic filter efficiently processes multiple queries, isolating only the structurally relevant pairs for down-stream geometric alignment.

### 4.3 Dual-Conformational Scanner

To accurately quantify the structural divergence masked by standard rigid-body superposition, we developed a dual-conformational scanner that models the protein as two independent rigid bodies. The algorithm utilizes the highly conserved SSE alignments generated from our previous framework [17]. To ensure that the resulting structural segments possess sufficient geometric complexity, the analysis is restricted to pairs sharing a minimum of six aligned SSEs.

The scanner systematically searches for the optimal structural hinge by iterating a cleavage point across the SSE alignment. To maintain the structural integrity of the resulting sub-domains, the sliding boundary is constrained to leave a minimum of three SSEs in both the N-terminal and C-terminal segments, an approach analogous to the self-scanning methodology utilized for internal symmetry detection [18].

At each candidate boundary, the atomic coordinates of both proteins are split into two independent sub-structures. The geometric similarity of these sub-structures is then individually evaluated using US-align. Crucially, to account for complex topological rearrangements such as domain swapping or circular permutations, the scanner calculates a length-weighted average TM-score for two distinct alignment scenarios:

1. **Direct Mapping:** Alignment of the corresponding sequential segments (i.e., N-term to N-term, and C-term to C-term).
2. **Cross Mapping:** Alignment of the swapped segments (i.e., N-term to C-term, and C-term to N-term).

The algorithm takes the maximum of these length-weighted TM-scores at each step[15]. The cleavage point that yields the highest overall adjusted TM-score is ultimately identified as the optimal domain boundary, and this maximum score is recorded as the true geometric similarity of the flexible pair.

### 4.4 Sequence Identity Calculation

Following the heuristic filtration and TM-score validation steps, the computational cost of evaluating the remaining conformationally flexible pairs was reduced to a highly manageable level. Consequently, pairwise sequence identities were directly computed for these candidates using the global alignment algorithm implemented in the Biopython package [27]. The alignments were performed employing the standard BLOSUM62 substitution matrix [28], configured with a gap-open penalty of 10 and a gap-extension penalty of 0.5.

### 4.5 Data Availability

The dataset containing the identified protein pairs and classified clusters of conformational changes is publicly available on Zenodo at https://doi.org/10.5281/zenodo.19236054.

## Notes

### Competing Interest Statement

The authors have declared no competing interest.

